# eTumorRisk, an algorithm predicts cancer risk based on comutated gene networks in an individual’s germline genome

**DOI:** 10.1101/393090

**Authors:** Jinfeng Zou, Edwin Wang

## Abstract

Early cancer detection has potentials to reduce cancer burden. A prior identification of the high-risk population of cancer will facilitate cancer early detection. Traditionally, cancer predisposition genes such as BRCA1/2 have been used for identifying high-risk population of developing breast and ovarian cancers. However, such high-risk genes have only a few. Moreover, the complexity of cancer hints multiple genes involved but also prevents from identifying such predictors for predicting high-risk subpopulation. Therefore, we asked if the germline genomes could be used to identify high-risk cancer population. So far, none of such predictive models has been developed. Here, by analyzing of the germline genomes of 3,090 cancer patients representing 12 common cancer types and 25,701 non-cancer individuals, we discovered significantly differential co-mutated gene pairs between cancer and non-cancer groups, and even between cancer types. Based on these findings, we developed a network-based algorithm, eTumorRisk, which enables to predict individuals’ cancer risk of six genetic-dominant cancers including breast, colon, brain, leukemia, ovarian and endometrial cancers with the prediction accuracies of 74.1-91.7% and have 1-3 false-negatives out of the validating samples (n=14,701). The eTumorRisk which has a very low false-negative rate might be useful in screening of general population for identifying high-risk cancer population.

## Introduction

Cancer is the leading cause of death in the world and the third largest burden in the healthcare system. Many cancers have been diagnosed at the middle-or late-stage (i.e., advanced cancer) at which time most tumors have spread and become incurable; therefore, improvements in overall survival and morbidity have been modest over the past few decades [1–5]. Historical data suggest that early detection of cancer is crucial for its ultimate control and prevention [6, 7]. Thus, it is envisioned that, besides the promotion of lifestyle changes, improving early diagnosis is the best strategy for reducing the impact of carcinogenesis. Ideally, screening of a subpopulation who has high-risk of developing cancer as early as possible could promote the early detection of cancer greatly. It has been reported that besides ‘hereditary’ cancers, many of ‘sporadic’ cancers have significant inherited component [8], suggesting the feasibility of identifying high-risk individuals based on germline genomes.

Traditionally, high-risk cancer predisposition genes (CPGs) have been used for identifying high-risk individuals. For example, Kuchenbaecker et al. reported that a woman who has BRCA1/2 mutations in her germline has a cumulative risk of 72%/69% (i.e., to the age of 80) of developing breast cancer and 44%/17% of ovarian cancer in her lifetime [9]. However, it is worth of noting that most of these BRCA1/2 carriers are involved in the study with family history concern. It was reported that patients with more family members having breast or ovarian cancers have higher frequencies of BRCA1/2 mutations than those with a single case [10]. It suggests that some unknown mechanisms promote the occurrence of BRCA1/2 mutations, which could help identify high-risk population. Similarly, 45% of the colon cancer cases are believed to be associated with a heritable factor, among which only 5-10% are related to the germline mutations in APC and DNA mismatch repair genes [11, 12]. To further understand cancer predispositions, studies have extended to moderate- and weak-risk CPGs [13–16], and the cumulative contributions of multiple germline mutations to cancer have been shown [17–20]. These findings suggest it is possible to identify high-risk people, however, so far it has been challenging to use germline mutations to predict who could bear cancer risk.

To meet this challenge, we analyzed 28,791 whole-exomes of germline genomes (3,090 cancer patients across 12 cancer types and 25,701 non-cancer individuals), and surprisingly found that a set of co-mutated gene pairs are preexisting in the germline genomes of cancer patients more frequently than in non-cancer germline genomes, and vice versa. Furthermore, each cancer type has its specifically co-mutated gene pairs. Based on these results, we developed a novel network-based algorithm, eTumorRisk, to identify high-risk individuals for cancers based on their germline genomic information. We validated the eTumorRisk in 14,701 non-cancer individuals and 1,098 cancer patients and showed its ability of identifying high-risk individuals for six genetic-dominant cancers (i.e., breast, colon, brain, leukemia, ovarian and endometrial cancers). The predictions of the eTumorRisk generated 0.0068% of false-positives in 14,701 non-cancer individuals. These results highlight that the eTumorRisk could be used for the screening of general population for selecting high-risk individuals of the six cancers to facilitate the early detection of cancer.

## Results

### Significantly differential co-mutated genes are encoded in the germline genomes between cancer and non-cancer groups, and even between cancer types

Although the germline genomes of several thousands of cancer patients have been sequenced, the differences between cancer patients’ germline genomes and non-cancer individuals’ have been rarely examined. To do so, we compiled the germline genomes of 3,090 cancer patients representing 12 common cancer types, and of 25,701 non-cancer individuals (Table 1). The functionally mutated genes were extracted from each genome sequence data (see Methods, all the mutated genes mentioned in the text are functionally mutated genes), and the frequencies of the mutated genes were evaluated in each cancer type. First, we compared the frequencies of the mutated genes between colon adenocarcinoma (COAD) and non-cancer individuals. We found that 518 genes were mutated significantly more frequently in COAD than in non-cancer individuals. Furthermore, 15,714 co-mutated gene pairs were significantly more frequent in COAD than in non-cancer individuals. Similar results were obtained when the analysis was extended to other cancer types. These results support a notion that the germline genomes of cancer patients have pre-existing mutated gene patterns which could functionally interact and contribute to cancer risk. To investigate whether each cancer type has its own specific set of co-mutated gene pairs, the comparison analyses between the 12 cancer types were performed in a pair-wise manner. Indeed, each cancer type does have its specific set. Taken together, these results suggest that cancer patients’ and non-cancer individuals’ germline genomes have distinct co-mutated gene pairs, and furthermore, each cancer type has its own specific co-mutated gene pairs. Clearly, these novel findings provide a solid foundation for developing a new algorithm for predicting of cancer risk based on co-mutated genes in germline genomes.

**Table 1.** Datasets and sample sizes.

| Sample | Abbreviation | No. of samples |
| --- | --- | --- |
| Cancer patients |  |  |
| Bladder urothelial carcinoma | BLCA | 212 |
| Breast invasive carcinoma (luminal subtype) | BRCA | 328 |
| Colon adenocarcinoma | COAD | 274 |
| Glioblastoma multiforme | GBM | 212 |
| Head and neck squamous cell carcinoma | HNSC | 227 |
| Kidney renal clear cell carcinoma | KIRC | 203 |
| Acute myeloid leukemia | LAML | 120 |
| Lung adenocarcinoma | LUAD | 348 |
| Lung squamous cell carcinoma | LUSC | 350 |
| Ovarian serous cystadenocarcinoma | OV | 244 |
| Skin cutaneous melanoma | SKCM | 359 |
| Uterine corpus endometrial carcinoma | UCEC | 213 |
| Non-cancer individuals |  |  |
| 1000 genome project | G1K | 2,431 |
| Non-cancer cohort | NC | 23,270 |

### Overview of the eTumorRisk algorithm

The eTumorRisk contains two components (Fig 1). The first component is to build network models for discriminating a cancer sample (of a cancer type) from non-cancer samples using co-mutated gene pairs in germline genomes. Because cancer types share some co-mutated gene pairs in germline genomes, a sample could be predicted to have a risk of developing multiple cancer types based on the first component, and therefore the second component is to determine which cancer type has the highest chance for the sample. Cancer heterogeneity is a major factor for impeding the robustness of predictive biomarkers. To address issue, we designed the eTumorRisk by (1) focusing on cancer hallmark genes to construct network models of discriminating a cancer type from non-cancer samples with the consideration of their mutations having higher chance to contribute to tumorigenesis than others. By doing so, we could filter the ‘noisy genes’ which are not related to tumorigenesis in germlines. (2) using the re-sampling technique to construct random networks and then cluster them into a few network groups and randomly select a network as representative for each group. The randomly resampled datasets were used to simulate populations, and the similar populations were clustered to extract the representative features of the networks. By doing so, we could gain more accurate and robust signals for predictions in both components. In addition, we used large number of the non-cancer samples for training (n=10,000) and validating (n=14,701) the algorithm, because the eTumorRisk was designed to screen the general population and the overall cancer incidence is 43.92 out of 10,000 samples per year (https://www.cancer.gov/about-cancer/understanding/statistics). By doing so, it might control false-positives.

**Fig. 1.**
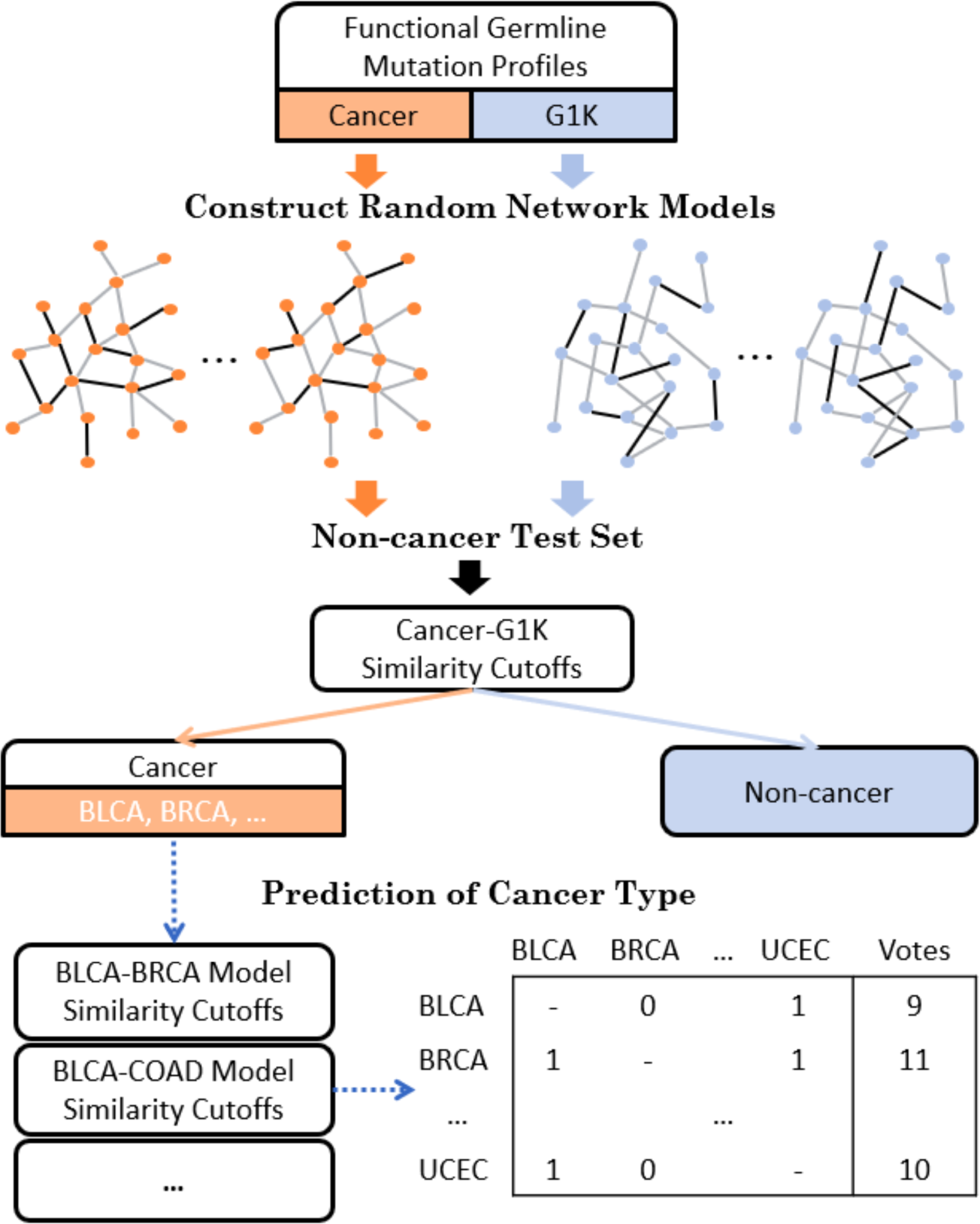
Prediction of cancer and specific cancer type. Each cancer and G1K (1000 genomes) functional germline mutation profiles were trained to construct random network models. Non-cancer test set was used as a control to select similarity cutoffs for prediction of cancer or non-cancer. Cancer candidates were predicted with a cancer type by the voting of all the pairs of cancer models.

The novelties of this algorithm are (1) constructing of germline co-mutated gene networks enabling to discriminate a cancer type from non-cancer samples, and between cancer types, (2) focusing on cancer molecular mechanisms (i.e., represented using cancer hallmarks) to filter out noise, (3) identifying multiple representative networks, each of which could represent a fraction of the samples within a cancer type (i.e., samples within a cancer type will be classified into several subgroups due to the heterogeneous nature of cancer), (4) controlling of false-positives by using large cohorts for training and validating the algorithm. Such large cohorts (n=25,701) have been rarely used for developing genome-based algorithms in the past.

### Cancer risk prediction and validation using the eTumorRisk

As mentioned above, we used 10,000 out of the 24,701 non-cancer samples to determine the similarity-cutoff for each cancer-determining predictive model, and then, we used the remaining 14,701 non-cancer samples to validate the similarity-cutoffs. Specifically, each similarity-cutoff for discriminating each cancer type from the non-cancer group was selected by controlling that none of the 10,000 non-cancer individuals was predicted to be a cancer sample. As shown in Table 2, none of the 14,701 non-cancer samples used for validation was predicted to be a cancer sample for breast invasive carcinoma (luminal subtype, BRCA), COAD, glioblastoma multiforme (GBM), acute myeloid leukemia (LAML) and ovarian serous cystadenocarcinoma (OV) cancer types. Similarly, 1 out of the 14,701 non-cancer samples was predicted to be head and neck squamous cell carcinoma (HNSC), kidney renal clear cell carcinoma (KIRC) and uterine corpus endometrial carcinoma (UCEC) cancer types based on the similarity cutoffs of their predictive models. These results suggest that the cutoffs of the predictive models for the cancer types mentioned above have been validated. As for bladder urothelial carcinoma (BLCA), lung adenocarcinoma (LUAD), lung squamous cell carcinoma (LUSC) and skin cutaneous melanoma (SKCM), the predictive models have not been established (i.e., the condition of selecting similarity-cutoff cannot be satisfied). Of note, BLCA, LUAD, LUSC and SKCM are well-known environmental-dominated cancers, whereas GBM, BRCA, COAD, LAML, OV and UCEC are genetic-dominated cancers. As shown in Table 2, for LAML, OV and BRCA, 75%, 74% and 61% were assigned to the correct cancer group, respectively, while for COAD, GBM, KIRC and UCEC, the percentages of the predictions reached ~40%. Taken together, we concluded that germline genetic variants could distinguish cancer-risk individuals from non-cancer individuals for the genetic-dominated cancers but not the environmental-dominated cancers.

**Table 2.** Cancer and cancer type prediction based on 10,000 non-cancer individuals.

| **Cancer prediction** |  |  |  | **Cancer type prediction** |  |  |
| --- | --- | --- | --- | --- | --- | --- |
| **Cancer** | **Non-cancer samples (n=14,701)** | **Cancer sample (n=1,098)** | **Power for cancer** | **No. of sample in the 1,098 samples** | **Accuracy** | **Power** |
| BLCA | - | - | 6 (9%)<sup>b</sup> | 64 | - | 0% |
| BRCA | 0<sup>a</sup> | 74<sup>a</sup> | 60 (61%) | 98 | 53.1% | 43.9% |
| COAD | 0 | 21 | 30 (37%) | 82 | 76% | 23.2% |
| GBM | 0 | 39 | 26 (41%) | 64 | 78.6% | 17.2% |
| HNSC | 1 | 21 | 9 (13%) | 68 | 0% | 0% |
| KIRC | 1 | 51 | 27 (44%) | 61 | 25% | 1.6% |
| LAML | 0 | 90 | 27 (75%) | 36 | 71.4% | 55.6% |
| LUAD | - | - | 13 (13%) | 104 | 16.7% | 0.9% |
| LUSC | - | - | 14 (13%) | 105 | 40% | 3.8% |
| OV | 0 | 84 | 54 (74%) | 73 | 91.5% | 58.9% |
| SKCM | - | - | 26 (24%) | 108 | 76.7% | 21.3% |
| UCEC | 1 | 110 | 24 (38%) | 64 | 25% | 3.1% |

<sup>a</sup>The number of samples predicated as cancer by each cancer-predictive model. For BRCA, OV and UCEC, only female patients predicted as cancer were considered. <sup>b</sup>The number of cancer samples predicted correctly as cancer by all the eight cancer-predictive models and the corresponding proportion in each cancer type shown in the parentheses. “-” represents not applicable because the false-positive control cannot be satisfied.

We found that some samples of a cancer type have been predicted to another cancer type or multiple cancer types. Overall, 53.1% of the predicted cancer samples were assigned into one cancer type, while 46.9% samples were predicted to be multiple cancer types. To further improve cancer type predictions, we applied cancer-type predictive models (i.e., the second component of the eTumorRisk) to assign the predicted cancer samples with one cancer type. Briefly, the pair-wised cancer predictive models were built, and the votes for each prediction were counted for deciding its cancer type (Methods). As shown in Table 2, the predictions of BRCA, COAD, GBM, LAML, OV, SKCM and UCEC cancer samples in the validation set reached higher prediction accuracies (i.e., the fraction of the correctly predicted samples of the total predicted samples) and better recall rates (i.e., the percentage of cancer patients of a cancer type who are correctly predicted as having the cancer). For example, for OV, the prediction accuracy and recall rate were 91.5% and 58.9%, respectively, while the accuracies and recall rates for COAD, GBM and SKCM were 76.0%-78.6% and 17.2% - 23.2%, respectively. As the eTumorRisk’s prediction is based on germline genomic information only, the recall rates of the predictions for most of the genetic-dominated cancers could not be very high. However, we found that the prediction recall rates for OV and LAML were as high as 58.9% and 55.6% %, respectively, suggesting that they may have high inheritability and can be captured by the eTumorRisk. In addition, we found that the samples of BRCA can be mistakenly assigned to UCEC, and vice versa. For example, 21 out of the 64 UCEC samples in the validation dataset were predicted to be BRCA samples, while 6 of the 98 BRCA samples were predicted to be UCEC samples (Supplementary Table 1). These results could be explained by the fact that BRCA and UCEC are two cancer types of woman-specific organs. When we combined these two cancer types together, the prediction accuracy and recall rate reached 80.9% and 44.4%, respectively. Based on these results, we suggested that if a woman is predicted to have high risk for either BRCA or UCEC, it is better to follow up that person for examining both cancer types. For the other cancer type predictions, none samples were predicted as BLCA, all the four samples predicted as HNSC were wrong predictions, while low accuracies and recall rates for KIRC, LUAD and LUSC (Table 2).

The reliability and applicability of the eTumorRisk were shown above. Therefore, we further evaluated the performance of eTumorRisk based on all the 24,701 non-cancer samples on that none of them are predicted wrongly, while only female non-cancer samples (8,169) were used for female-related cancer types (BRCA, OV and UCEC). The powers for cancer predictions decreased a little and the accuracies remained similar levels for six genetic-dominant cancers and for most of the other cancer types (Table 3). Specifically, the accuracy of the COAD predictions increased from 0.76 to 0.86 because the three wrongly assigned samples were not identified as cancer by the first component of eTumorRisk (Supplementary Table 1 and 2), while the recall rate was sustained as 23.2%, suggesting the stability of COAD prediction by eTumorRisk. Similar result was for LAML, while the power decreased only ~3% for GBM and SKCM. The accuracy and power for the united BRCA and UCEC predictions were 79.1% and 33% (Supplementary Table 2), showing stable accuracy after involving more female non-cancer samples.

**Table 3.** Cancer and cancer type prediction based on 24,701 non-cancer samples.

| Cancer | No. of sample in the 1,098 samples | Cancer prediction | Cancer type prediction |  |
| --- | --- | --- | --- | --- |
|  |  | Power for cancer | Accuracy | Power |
| BLCA | 64 | 5 (8%) | - | 0% |
| BRCA<sup>a</sup> | 98 | 45 (46%) | 53.1% | 34.7% |
| COAD | 82 | 27 (33%) | 86.4% | 23.2% |
| GBM | 64 | 26 (41%) | 75% | 14.1% |
| HNSC | 68 | 4 (6%) | 0% | 0% |
| KIRC | 61 | 20 (33%) | 0% | 0% |
| LAML | 36 | 27 (75%) | 74.1% | 55.6% |
| LUAD | 104 | 10 (10%) | 0% | 0% |
| LUSC | 105 | 11 (10%) | 42.9% | 2.9% |
| OV<sup>a</sup> | 73 | 55 (75%) | 91.7% | 60.3% |
| SKCM | 108 | 21 (19%) | 76% | 17.6% |
| UCEC<sup>a</sup> | 64 | 18 (28%) | 33% | 1.6% |

<sup>a</sup> For BRCA, OV and UCEC, only female cancer and non-cancer patients were considered.

## Discussion

In this study, we conducted the analysis of mutated genes of 3,090 cancer patients’ germline genomes and 25,701 non-cancer individuals’ genomes. We showed that distinct co-mutated gene pairs have been encoded in the germline genomes of the cancer patients from distinct cancer types by comparing the co-occurrence between a cancer and non-cancer groups and cancer types. These results support our previous hypothesis that pre-existing germline mutations could determine cancer risk and evolutionarily select biological pathways of tumors [21]. Based on these results, we developed the eTumorRisk to predict cancer risk of 12 cancer types using germline genomic information. Based on the predictions in ~14,701 non-cancer samples and 1,098 cancer samples, eTumorRisk achieved prediction accuracies of >74% for COAD, GBM, LAML, and SKCM. Based on the predictions in 8,169 female non-cancer samples and 235 OV, BRCA and UCEC samples, eTumorRisk achieved prediction accuracies of 0.92 and 0.79 for OV and the combination of BRCA and UCEC, respectively. However, eTumorRisk didn’t achieve good prediction performances in other cancer types of the 12 cancer types examined. Because eTumorRisk is a prediction tool based on purely germline genomics, which could be not suitable for predicting the risk of the cancer types which are induced predominantly by environmental and lifestyle factors. For example, lung and bladder cancers are well-known cancers which are induced by tobacco smoking (i.e., >85% of lung cancer patients are tobacco smokers). It is worth of noting that SKCM is believed to be environment-dominant cancer, however, there is a part of SKCM can be predicted by the non-SKCM predictive models in the first component and the second component of the eTumorRisk. It suggests that there are multiple subtypes existing in SKCM. Some studies have shown the predisposition of SKCM [22, 23]. On the other hand, eTumorRisk is developed for predicting the risk of the cancer types which are genetics-dominant such as BRCA, COAD, GBM, KIRC, LAML, OV and UCEC.

It has been proposed that early detection of cancer patients could save millions of lives for cancer patients. Therefore, with the advance of genome technology, cancer early detection has become an active area of research in the past few years. The advantages of cancer risk diagnosis will facilitate cancer early detection by identifying a high-risk subpopulation and monitoring their cancer development. In addition, prevention strategies could be applied to the high-risk population. For example, bilateral prophylactic mastectomy can reduce the risk of developing breast cancer by > 95% in women with BRCA1/2 mutations and by up to 90% in women who have a strong family history ([24, 25] and https://www.cancer.gov/types/breast/risk-reducing-surgery-fact-sheet). Drug tamoxifen is shown to prevent the development of breast cancer in healthy women who are determined at increased risk of developing breast cancer [26]; anastrozole and metformin are also to help prevent cancer development [27–29]. Besides, the inflammatory process is shown to be one of the predisposing conditions for the initiation and development of tumor [30]. The extrinsic inflammation can be caused by autoimmune diseases, obesity, smoking, asbestos exposure and alcohol, while the intrinsic inflammation can be triggered by mutations [30]. Both inflammation can induce immune suppression, and thereby promote the initiation and progression of tumor. It has been proven that lifestyle modifications and the anti-inflammatory drugs (e.g. aspirin) can significantly reduce cancer risk [30, 31]. We believe these will help alleviate the development of cancer, and even possibly avoid cancer occurrence.

## Materials and Methods

### Sequencing data pre-processing and variant calling

Twelve cancer types from The Cancer Genome Atlas (TCGA) project [32] were analyzed in this study (Table 1). The whole-exome sequencing data were downloaded from the data portal, and processed as described previously [21]. Briefly, low quality reads were filtered by the picard-tools, and duplicates were also marked up by the same tool (http://broadinstitute.github.io/picard). Reads with low mapping quality (<=60) were filtered by BamTools [33]. Local realignment around indels and base recalibration were done by the GATK tool [34]. Variant calling was done by Varscan 2 [35].

The non-cancer samples were from a collected cohort including samples of the 1000 Genome project [36]. All samples of the 1000 Genome project were downloaded, pre-processed with the procedure shown above and the variant calling was also done by Varscan 2. For the other samples, the variant matrices were downloaded from the resources.

### Germline mutation extraction and functional annotation

Germline mutations were defined reliably by controlling variant allele frequency (VAF) no less than 90% for a homozygous mutation and between 45% and 55% for a heterozygous mutation in normal (blood) samples. The functional annotation of germline mutations was evaluated on the effect of sequence changes on proteins using Sorting Intolerant From Tolerant (SIFT), PolyPhen [37] and MutationTaster2 [38]. A mutation is defined as functional if one of the three algorithms predicts it as function-related. Next, these functional germline mutations were summarized on gene level.

### Training, test, and reference sets

The training sets included 12 cancer types and healthy people from the 1000 Genome project (G1K). For cancers, 70% samples randomly sampling from the whole set were used as the training set and the left 30% were as the test set. For non-cancer group, the randomly sampled 1000 samples from the 1000 genome project were used as the training set. The remaining 1,431 samples of G1K and the other non-cancer cohorts were remained as a reference set to select similarity score (see below) for determining cancer or non-cancer group for the cancer test sets (see details in Results).

### Cancer hallmark-associated genes and protein-protein interaction (PPI) network

The cancer hallmark-associated genes were collected through Gene Ontology (GO) annotation [39] based on cancer hallmarks [40], and two publications about cancer genes [41, 42]. To incorporate more related genes, the annotated gene list was extended by including genes having at least three links to cancer hallmark-associated genes on the PPI network which is composed of 12,612 genes and 175,696 edges.

### Selection of differentially mutated genes, differential co-occurrences and generation of mutation network templates

Differentially mutated genes (DMG) were selected by evaluating the mutation frequency differences of the preselected cancer hallmark-associated genes between cancer and non-cancer groups based on their z-score transformation (*P*<0.05). Then, the differential co-occurrences among the selected DMGs between cancer and non-cancer groups were identified based on 1000 random experiments in each group by randomly shuffling samples for each gene (*P*<0.05). The selected significant co-mutations generated the mutation network templates for cancer and non-cancer groups or between cancer types. We proposed to use the bilateral networks produced by comparing cancer to the non-cancer group, or one cancer type to another, and vice versa, to develop the eTumorRisk. We tended to build the bilateral template networks which include all possible truly discriminative co-mutated gene pairs given by the maximization of the sample size (i.e., all the training samples). The significant co-mutations in the cancer network template have higher frequency than their counterparts in the normal network template, and vice versa. The same methods were applied to identify DMGs between each pair of cancer types, and thereby build the mutation network templates.

### Heterogeneous mutation network models

To deal with the cancer heterogeneity, a model panel was generated to capture the heterogeneity by two steps. First, a sufficiently large number of random datasets (*e.g.* 2500) were sampled from the training set by 50% to get sub-population mutation network models for each pair of comparison (*i.e.* each cancer against the non-cancer group, or between cancer types). Then, the models were mapped to the corresponding network templates (i.e. cancer or non-cancer), and the links which have high mutation frequency (*e.g.* 1%) and frequency ratio between the comparison (*e.g.* 2) were retained to generate the final network model for achieving high efficiency. The clustering of these sub-population network models was done for capturing the diversity within each group (cluster R package). The number of clusters was selected based on a silhouette evaluation upon 1000 possible classifications (*i.e.* 1, 2, …, 1000 clusters). For example, there were 56 network subgroups classified for the random networks of the COAD group, while 50 subgroups selected for the non-cancer group. Thereafter, a representative sub-population network model was randomly selected from each cluster and used for the subsequent analysis. Summarily, two sets of representative sub-population network models were generated for the pair of comparison. To save computing time in the process of clustering the large sub-population networks, each network (i.e. a matrix) was first shrunk by grouping the adjacent ten genes as a sub-network and then assigning it with the counting value of the links in this sub-network.

### Prediction of cancer or non-cancer group

For a sample, its similarity to each representative sub-population network model was evaluated by the Jaccard index between its mutation network and the network model (the number of overlapped links divided by the number of union of links). Then, the Jaccard scores were clustered using PAM (Partitioning Around Medoids, pamr R package) combined with Silhouette index within groups (e.g. cancer and non-cancer). The clustering was controlled to a certain number of clusters (*e.g.* no larger than 10) for the balance of reflecting the diversity of heterogeneity and achieving robustness. After classification, the average Jaccard score was calculated for each cluster and then the maximum Jaccard score was used. The operation of the clustering is for promoting robustness and the use of maximum score is for identifying the most possible network model that the sample could be. Then, the ratio of Jaccard scores between cancer and non-cancer groups or vice versa was calculated as the similarity score to cancer or non-cancer group.

To assign a sample to cancer or non-cancer group, a similarity cutoff was selected based on the predictions of the non-cancer dataset. First, 10,000 samples of the non-cancer dataset were used to get a similarity cutoff which make none of them predicted as cancer. The large training set is used to control false-positives. Then, it was tested on the remaining 14,701 non-cancer samples. Finally, the similarity score was used to predict cancer or non-cancer for the cancer test dataset.

### Prediction of cancer type

Similar procedure for assigning a sample to cancer and non-cancer group (shown above) was applied to discriminate cancer types for samples predicted into cancer group. But, the ratio cutoffs for a pair of cancer types were selected using OptimalCutpoints R package based on the training set [43]. A sample is screened by the network models of all pair-wised cancer types. The prediction scores were summarized for each possible cancer type and ranked for the final decision (Fig. 1).

---

**Table S1.** Distribution of predicted cancer samples based on 10,000 non-cancer individuals.

| **Cancer** | **No. of samples predicted<sup>a</sup>** | **BLCA** | **BRCA** | **COAD** | **GBM** | **HNSC** | **KIRC** | **LAML** | **LUAD** | **LUSC** | **OV** | **SKCM** | **UCEC** |
| --- | --- | --- | --- | --- | --- | --- | --- | --- | --- | --- | --- | --- | --- |
| BLCA | 0 | 0 | 0 | 0 | 0 | 0 | 0 | 0 | 0 | 0 | 0 | 0 | 0 |
| BRCA | 81 | 0 | 43 | 3 | 0 | 0 | 13 | 0 | 0 | 0 | 1 | 0 | 21 |
| COAD | 25 | 0 | 2 | 19 | 0 | 0 | 0 | 0 | 4 | 0 | 0 | 0 | 0 |
| GBM | 14 | 0 | 0 | 0 | 11 | 0 | 0 | 1 | 0 | 1 | 1 | 0 | 0 |
| HNSC | 2 | 0 | 0 | 0 | 0 | 0 | 0 | 0 | 2 | 0 | 0 | 0 | 0 |
| KIRC | 4 | 0 | 0 | 0 | 0 | 0 | 1 | 0 | 0 | 1 | 1 | 1 | 0 |
| LAML | 28 | 0 | 0 | 0 | 5 | 0 | 0 | 20 | 0 | 0 | 3 | 0 | 0 |
| LUAD | 6 | 4 | 0 | 0 | 0 | 0 | 0 | 0 | 1 | 1 | 0 | 0 | 0 |
| LUSC | 10 | 1 | 0 | 0 | 0 | 2 | 1 | 0 | 2 | 4 | 0 | 0 | 0 |
| OV | 47 | 0 | 0 | 0 | 2 | 0 | 0 | 2 | 0 | 0 | 43 | 0 | 0 |
| SKCM | 30 | 0 | 0 | 0 | 0 | 2 | 0 | 0 | 1 | 4 | 0 | 23 | 0 |
| UCEC | 8 | 0 | 6 | 0 | 0 | 0 | 0 | 0 | 0 | 0 | 0 | 0 | 2 |

<sup>a</sup> The number of samples predicted as a cancer type. <sup>b</sup> The number of samples belonging to each cancer type (listed as columns) in the predicted samples for the given cancer type.

**Table S2.** Distribution of predicted cancer samples based on 24,701 non-cancer individuals.

| **Cancer** | **No. of samples predicted<sup>a</sup>** | **BLCA** | **BRCA** | **COAD** | **GBM** | **HNSC** | **KIRC** | **LAML** | **LUAD** | **LUSC** | **OV** | **SKCM** | **UCEC** |
| --- | --- | --- | --- | --- | --- | --- | --- | --- | --- | --- | --- | --- | --- |
| BLCA | 0 | 0 | 0 | 0 | 0 | 0 | 0 | 0 | 0 | 0 | 0 | 0 | 0 |
| BRCA | 64 | 0 | 34 | 3 | 0 | 0 | 10 | 0 | 0 | 0 | 1 | 0 | 16 |
| COAD | 22 | 0 | 1 | 19 | 0 | 0 | 0 | 0 | 2 | 0 | 0 | 0 | 0 |
| GBM | 12 | 0 | 0 | 0 | 9 | 0 | 0 | 1 | 0 | 1 | 1 | 0 | 0 |
| HNSC | 1 | 0 | 0 | 0 | 0 | 0 | 0 | 0 | 1 | 0 | 0 | 0 | 0 |
| KIRC | 2 | 0 | 0 | 0 | 0 | 0 | 0 | 0 | 0 | 1 | 1 | 0 | 0 |
| LAML | 27 | 0 | 0 | 0 | 4 | 0 | 0 | 20 | 0 | 0 | 3 | 0 | 0 |
| LUAD | 3 | 3 | 0 | 0 | 0 | 0 | 0 | 0 | 0 | 0 | 0 | 0 | 0 |
| LUSC | 7 | 1 | 0 | 0 | 0 | 2 | 0 | 0 | 1 | 3 | 0 | 0 | 0 |
| OV | 48 | 0 | 0 | 0 | 2 | 0 | 0 | 2 | 0 | 0 | 44 | 0 | 0 |
| SKCM | 25 | 0 | 0 | 0 | 0 | 1 | 0 | 0 | 1 | 4 | 0 | 19 | 0 |
| UCEC | 3 | 0 | 2 | 0 | 0 | 0 | 0 | 0 | 0 | 0 | 0 | 0 | 1 |

<sup>a</sup> The number of samples predicted as a cancer type. <sup>b</sup> The number of samples belonging to each cancer type (listed as columns) in the predicted samples for the given cancer type.

## Reference

[1] Aravanis, A.M.; Lee, M.; Klausner, R.D. Next-Generation Sequencing of Circulating Tumor DNA for Early Cancer Detection. Cell, 2017, 168(4), 571-574.

[2] Cho, H.; Mariotto, A.B.; Schwartz, L.M.; Luo, J.; Woloshin, S. When do changes in cancer survival mean progress? The insight from population incidence and mortality. J Natl Cancer Inst Monogr, 2014, 2014(49), 187-197.

[3] Sakhuja, S.; Yun, H.; Pisu, M.; Akinyemiju, T. Availability of healthcare resources and epithelial ovarian cancer stage of diagnosis and mortality among Blacks and Whites. J Ovarian Res, 2017, 10(1), 57.

[4] Lin, Y.; Wimberly, M.C. Geographic Variations of Colorectal and Breast Cancer Late-Stage Diagnosis and the Effects of Neighborhood-Level Factors. J Rural Health, 2017, 33(2), 146-157.

[5] Temkin, S.M.; Terplan, M. Trends in Relative Survival for Ovarian Cancer From 1975-2011. Obstet Gynecol, 2015, 126(4), 898.

[6] Hiom, S.C. Diagnosing cancer earlier: reviewing the evidence for improving cancer survival. Br J Cancer, 2015, 112 Suppl 1(S1-5.

[7] Butler, T.M.; Spellman, P.T.; Gray, J. Circulating-tumor DNA as an early detection and diagnostic tool. Curr Opin Genet Dev, 2017, 42(14-21.

[8] Lu, Y.; Ek, W.E.; Whiteman, D.; Vaughan, T.L.; Spurdle, A.B.; Easton, D.F.; Pharoah, P.D.; Thompson, D.J.; Dunning, A.M.; Hayward, N.K.; Chenevix-Trench, G.; Macgregor, S. Most common 'sporadic' cancers have a significant germline genetic component. Hum Mol Genet, 2014, 23(22), 6112-6118.

[9] Kuchenbaecker, K.B.; Hopper, J.L.; Barnes, D.R.; Phillips, K.A.; Mooij, T.M.; Roos-Blom, M.J.; Jervis, S.; van Leeuwen, F.E.; Milne, R.L.; Andrieu, N.; Goldgar, D.E.; Terry, M.B.; Rookus, M.A.; Easton, D.F.; Antoniou, A.C.; McGuffog, L.; Evans, D.G.; Barrowdale, D.; Frost, D.; Adlard, J.; Ong, K.R.; Izatt, L.; Tischkowitz, M.; Eeles, R.; Davidson, R.; Hodgson, S.; Ellis, S.; Nogues, C.; Lasset, C.; Stoppa-Lyonnet, D.; Fricker, J.P.; Faivre, L.; Berthet, P.; Hooning, M.J.; van der Kolk, L.E.; Kets, C.M.; Adank, M.A.; John, E.M.; Chung, W.K.; Andrulis, I.L.; Southey, M.; Daly, M.B.; Buys, S.S.; Osorio, A.; Engel, C.; Kast, K.; Schmutzler, R.K.; Caldes, T.; Jakubowska, A.; Simard, J.; Friedlander, M.L.; McLachlan, S.A.; Machackova, E.; Foretova, L.; Tan, Y.Y.; Singer, C.F.; Olah, E.; Gerdes, A.M.; Arver, B.; Olsson, H. Risks of Breast, Ovarian, and Contralateral Breast Cancer for BRCA1 and BRCA2 Mutation Carriers. JAMA, 2017, 317(23), 2402-2416.

[10] Kast, K.; Rhiem, K.; Wappenschmidt, B.; Hahnen, E.; Hauke, J.; Bluemcke, B.; Zarghooni, V.; Herold, N.; Ditsch, N.; Kiechle, M.; Braun, M.; Fischer, C.; Dikow, N.; Schott, S.; Rahner, N.; Niederacher, D.; Fehm, T.; Gehrig, A.; Mueller-Reible, C.; Arnold, N.; Maass, N.; Borck, G.; de Gregorio, N.; Scholz, C.; Auber, B.; Varon-Manteeva, R.; Speiser, D.; Horvath, J.; Lichey, N.; Wimberger, P.; Stark, S.; Faust, U.; Weber, B.H.; Emons, G.; Zachariae, S.; Meindl, A.; Schmutzler, R.K.; Engel, C. Prevalence of BRCA1/2 germline mutations in 21 401 families with breast and ovarian cancer. J Med Genet, 2016, 53(7), 465-471.

[11] Hahn, M.M.; de Voer, R.M.; Hoogerbrugge, N.; Ligtenberg, M.J.; Kuiper, R.P.; van Kessel, A.G. The genetic heterogeneity of colorectal cancer predisposition - guidelines for gene discovery. Cell Oncol (Dordr), 2016, 39(6), 491-510.

[12] de Voer, R.M.; Hahn, M.M.; Weren, R.D.; Mensenkamp, A.R.; Gilissen, C.; van Zelst-Stams, W.A.; Spruijt, L.; Kets, C.M.; Zhang, J.; Venselaar, H.; Vreede, L.; Schubert, N.; Tychon, M.; Derks, R.; Schackert, H.K.; Geurts van Kessel, A.; Hoogerbrugge, N.; Ligtenberg, M.J.; Kuiper, R.P. Identification of Novel Candidate Genes for Early-Onset Colorectal Cancer Susceptibility. PLoS Genet, 2016, 12(2), e1005880.

[13] Sokolenko, A.P.; Preobrazhenskaya, E.V.; Aleksakhina, S.N.; Iyevleva, A.G.; Mitiushkina, N.V.; Zaitseva, O.A.; Yatsuk, O.S.; Tiurin, V.I.; Strelkova, T.N.; Togo, A.V.; Imyanitov, E.N. Candidate gene analysis of BRCA1/2 mutation-negative high-risk Russian breast cancer patients. Cancer Lett, 2015, 359(2), 259-261.

[14] Han, M.R.; Zheng, W.; Cai, Q.; Gao, Y.T.; Zheng, Y.; Bolla, M.K.; Michailidou, K.; Dennis, J.; Wang, Q.; Dunning, A.M.; Brennan, P.; Chen, S.T.; Choi, J.Y.; Hartman, M.; Ito, H.; Lophatananon, A.; Matsuo, K.; Miao, H.; Muir, K.; Sangrajrang, S.; Shen, C.Y.; Teo, S.H.; Tseng, C.C.; Wu, A.H.; Yip, C.H.; Kang, D.; Xiang, Y.B.; Easton, D.F.; Shu, X.O.; Long, J. Evaluating genetic variants associated with breast cancer risk in high and moderate-penetrance genes in Asians. Carcinogenesis, 2017, 38(5), 511-518.

[15] Maxwell, K.N.; Wubbenhorst, B.; D'Andrea, K.; Garman, B.; Long, J.M.; Powers, J.; Rathbun, K.; Stopfer, J.E.; Zhu, J.; Bradbury, A.R.; Simon, M.S.; DeMichele, A.; Domchek, S.M.; Nathanson, K.L. Prevalence of mutations in a panel of breast cancer susceptibility genes in BRCA1/2-negative patients with early-onset breast cancer. Genet Med, 2015, 17(8), 630-638.

[16] Song, H.; Dicks, E.; Ramus, S.J.; Tyrer, J.P.; Intermaggio, M.P.; Hayward, J.; Edlund, C.K.; Conti, D.; Harrington, P.; Fraser, L.; Philpott, S.; Anderson, C.; Rosenthal, A.; Gentry-Maharaj, A.; Bowtell, D.D.; Alsop, K.; Cicek, M.S.; Cunningham, J.M.; Fridley, B.L.; Alsop, J.; Jimenez-Linan, M.; Hogdall, E.; Hogdall, C.K.; Jensen, A.; Kjaer, S.K.; Lubinski, J.; Huzarski, T.; Jakubowska, A.; Gronwald, J.; Poblete, S.; Lele, S.; Sucheston-Campbell, L.; Moysich, K.B.; Odunsi, K.; Goode, E.L.; Menon, U.; Jacobs, I.J.; Gayther, S.A.; Pharoah, P.D. Contribution of Germline Mutations in the RAD51B, RAD51C, and RAD51D Genes to Ovarian Cancer in the Population. J Clin Oncol, 2015, 33(26), 2901-2907.

[17] Crawford, B.; Adams, S.B.; Sittler, T.; van den Akker, J.; Chan, S.; Leitner, O.; Ryan, L.; Gil, E.; van't Veer, L. Multi-gene panel testing for hereditary cancer predisposition in unsolved high-risk breast and ovarian cancer patients. Breast Cancer Res Treat, 2017, 163(2), 383-390.

[18] Hildebrandt, M.A.; Reyes, M.E.; Lin, M.; He, Y.; Nguyen, S.V.; Hawk, E.T.; Wu, X. Germline Genetic Variants in the Wnt/beta-Catenin Pathway as Predictors of Colorectal Cancer Risk. Cancer Epidemiol Biomarkers Prev, 2016, 25(3), 540-546.

[19] Aggarwal, N.; Donald, N.D.; Malik, S.; Selvendran, S.S.; McPhail, M.J.; Monahan, K.J. The Association of Low-Penetrance Variants in DNA Repair Genes with Colorectal Cancer: A Systematic Review and Meta-Analysis. Clin Transl Gastroenterol, 2017, 8(7), e109.

[20] Shieh, Y.; Eklund, M.; Madlensky, L.; Sawyer, S.D.; Thompson, C.K.; Stover Fiscalini, A.; Ziv, E.; Van't Veer, L.J.; Esserman, L.J.; Tice, J.A. Breast Cancer Screening in the Precision Medicine Era: Risk-Based Screening in a Population-Based Trial. J Natl Cancer Inst, 2017, 109(5).

[21] Zaman, N.; Li, L.; Jaramillo, M.L.; Sun, Z.; Tibiche, C.; Banville, M.; Collins, C.; Trifiro, M.; Paliouras, M.; Nantel, A.; O'Connor-McCourt, M.; Wang, E. Signaling network assessment of mutations and copy number variations predict breast cancer subtype-specific drug targets. Cell Rep, 2013, 5(1), 216-223.

[22] Ward, K.A.; Lazovich, D.; Hordinsky, M.K. Germline melanoma susceptibility and prognostic genes: a review of the literature. J Am Acad Dermatol, 2012, 67(5), 1055-1067.

[23] Udayakumar, D.; Mahato, B.; Gabree, M.; Tsao, H. Genetic determinants of cutaneous melanoma predisposition. Semin Cutan Med Surg, 2010, 29(3), 190-195.

[24] Metcalfe, K.; Gershman, S.; Ghadirian, P.; Lynch, H.T.; Snyder, C.; Tung, N.; Kim-Sing, C.; Eisen, A.; Foulkes, W.D.; Rosen, B.; Sun, P.; Narod, S.A. Contralateral mastectomy and survival after breast cancer in carriers of BRCA1 and BRCA2 mutations: retrospective analysis. BMJ, 2014, 348(g226.

[25] Meijers-Heijboer, H.; van Geel, B.; van Putten, W.L.; Henzen-Logmans, S.C.; Seynaeve, C.; Menke-Pluymers, M.B.; Bartels, C.C.; Verhoog, L.C.; van den Ouweland, A.M.; Niermeijer, M.F.; Brekelmans, C.T.; Klijn, J.G. Breast cancer after prophylactic bilateral mastectomy in women with a BRCA1 or BRCA2 mutation. N Engl J Med, 2001, 345(3), 159-164.

[26] Cuzick, J.; Sestak, I.; Cawthorn, S.; Hamed, H.; Holli, K.; Howell, A.; Forbes, J.F. Tamoxifen for prevention of breast cancer: extended long-term follow-up of the IBIS-I breast cancer prevention trial. Lancet Oncol, 2015, 16(1), 67-75.

[27] Kasznicki, J.; Sliwinska, A.; Drzewoski, J. Metformin in cancer prevention and therapy. Ann Transl Med, 2014, 2(6), 57.

[28] Cuzick, J.; Sestak, I.; Forbes, J.F.; Dowsett, M.; Knox, J.; Cawthorn, S.; Saunders, C.; Roche, N.; Mansel, R.E.; von Minckwitz, G.; Bonanni, B.; Palva, T.; Howell, A. Anastrozole for prevention of breast cancer in high-risk postmenopausal women (IBIS-II): an international, double-blind, randomised placebo-controlled trial. Lancet, 2014, 383(9922), 1041-1048.

[29] Zi, F.; Zi, H.; Li, Y.; He, J.; Shi, Q.; Cai, Z. Metformin and cancer: An existing drug for cancer prevention and therapy. Oncol Lett, 2018, 15(1), 683-690.

[30] Todoric, J.; Antonucci, L.; Karin, M. Targeting Inflammation in Cancer Prevention and Therapy. Cancer Prev Res (Phila), 2016, 9(12), 895-905.

[31] Thompson, R. Preventing cancer: the role of food, nutrition and physical activity. J Fam Health Care, 2010, 20(3), 100-102.

[32] Cancer Genome Atlas Network. Comprehensive molecular portraits of human breast tumours. Nature, 2012, 490(7418), 61-70.

[33] Barnett, D.W.; Garrison, E.K.; Quinlan, A.R.; Stromberg, M.P.; Marth, G.T. BamTools: a C++ API and toolkit for analyzing and managing BAM files. Bioinformatics, 2011, 27(12), 1691-1692.

[34] DePristo, M.A.; Banks, E.; Poplin, R.; Garimella, K.V.; Maguire, J.R.; Hartl, C.; Philippakis, A.A.; del Angel, G.; Rivas, M.A.; Hanna, M.; McKenna, A.; Fennell, T.J.; Kernytsky, A.M.; Sivachenko, A.Y.; Cibulskis, K.; Gabriel, S.B.; Altshuler, D.; Daly, M.J. A framework for variation discovery and genotyping using next-generation DNA sequencing data. Nat Genet, 2011, 43(5), 491-498.

[35] Koboldt, D.C.; Zhang, Q.; Larson, D.E.; Shen, D.; McLellan, M.D.; Lin, L.; Miller, C.A.; Mardis, E.R.; Ding, L.; Wilson, R.K. VarScan 2: somatic mutation and copy number alteration discovery in cancer by exome sequencing. Genome Res, 2012, 22(3), 568-576.

[36] Auton, A.; Brooks, L.D.; Durbin, R.M.; Garrison, E.P.; Kang, H.M.; Korbel, J.O.; Marchini, J.L.; McCarthy, S.; McVean, G.A.; Abecasis, G.R. A global reference for human genetic variation. Nature, 2015, 526(7571), 68-74.

[37] Kumar, P.; Henikoff, S.; Ng, P.C. Predicting the effects of coding non-synonymous variants on protein function using the SIFT algorithm. Nat Protoc, 2009, 4(7), 1073-1081.

[38] Schwarz, J.M.; Cooper, D.N.; Schuelke, M.; Seelow, D. MutationTaster2: mutation prediction for the deep-sequencing age. Nat Methods, 2014, 11(4), 361-362.

[39] Gene Ontology Consortium. Gene Ontology Consortium: going forward. Nucleic Acids Res, 2015, 43(Database issue), D1049-1056.

[40] Hanahan, D.; Weinberg, R.A. The hallmarks of cancer. Cell, 2000, 100(1), 57-70.

[41] Futreal, P.A.; Coin, L.; Marshall, M.; Down, T.; Hubbard, T.; Wooster, R.; Rahman, N.; Stratton, M.R. A census of human cancer genes. Nat Rev Cancer, 2004, 4(3), 177-183.

[42] Rahman, N. Realizing the promise of cancer predisposition genes. Nature, 2014, 505(7483), 302-308.

[43] López-Ratón, M.; Rodríguez-Álvarez, M.X.; Suárez, C.C.; Sampedro, F.G. OptimalCutpoints: an R package for selecting optimal cutpoints in diagnostic tests. J Stat Softw, 2014, 61( ), 1-36.

